# Artificial intelligence-driven morphology-based enrichment of malignant cells from body fluid

**DOI:** 10.1101/2023.01.24.525423

**Authors:** Anastasia Mavropoulos, Chassidy Johnson, Vivian Lu, Jordan Nieto, Emilie Schneider, Kiran Saini, Michael L. Phelan, Linda Hsie, Maggie Wang, Janifer Cruz, Jeanette Mei, Julie Kim, Zhouyang Lian, Nianzhen Li, Stephane C. Boutet, Amy Wong-Thai, Weibo Yu, Qing-Yi Lu, Teresa Kim, Yipeng Geng, Maddison (Mahdokht) Masaeli, Thomas D. Lee, Jianyu Rao

**Affiliations:** Deepcell, Inc., 4025 Bohannon Dr., Menlo Park, CA, 94025; Pathology and Laboratory Medicine, University of California Los Angeles (UCLA), Los Angeles, CA, 90095

**Keywords:** morphology, cytology, effusion, carcinoma, high-throughput, machine learning, deep learning, image analysis, enrichment, computer vision

## Abstract

Cell morphology is a fundamental feature used to evaluate patient specimens in pathological analysis. However, traditional cytopathology analysis of patient effusion samples is limited by low tumor cell abundance coupled with high background of non-malignant cells, restricting the ability for downstream molecular and functional analyses to identify actionable therapeutic targets. We applied the Deepcell platform that combines microfluidic sorting, brightfield imaging, and real-time deep learning interpretations based on multi-dimensional morphology to enrich carcinoma cells from malignant effusions without cell staining or labels. Carcinoma cell enrichment was validated with whole genome sequencing and targeted mutation analysis, which showed higher sensitivity for detection of tumor fractions and critical somatic variant mutations that were initially at low-levels or undetectable in pre-sort patient samples. Combined, our study demonstrates the feasibility and added value of supplementing traditional morphology-based cytology with deep learning, multi-dimensional morphology analysis, and microfluidic sorting.

## INTRODUCTION

Cell morphology is a fundamental cell feature associated with cell identity, state, function, and disease. In recent reports, cell morphology has been used as key measurements in diverse applications including small molecule screening in cancer and infectious diseases, evaluating drug resistance, cell profiling to infer mechanism of action, and assessing metastatic capacity ^1–7^. With the advent of quantitative single cell imaging technology, morphology profiling has emerged as a fundamental readout to define the functional states, such as metastatic potential, of individual cells ^7^. While strides have been made in diverse applications to utilize quantitative profiling to systematically define morphological features, pathology workflows still largely rely on manual morphological observations. Trained pathologists analyze malignant cells in patient samples based on cell morphology, with the aid of dyes/stains, but these assessments may vary depending on individual pathologists and yield limited quantitative data, thereby limiting standardized information derived from tumor specimens.

Importantly, these traditional processes are unable to detect specific molecular targets that are crucial for precision care. With the availability of targeted therapies for certain genetic alterations, molecular testing has now become standard of care in cancer treatment ^8,9^. For instance, tyrosine kinase inhibitors are currently used to treat lung cancer in the presence of certain mutations ^10^, while tumors with high mutational burden (TMB) are responsive to immune checkpoint inhibitors ^11^. Thus, in the era of precision medicine, detecting specific actionable molecular targets is more important than merely confirming the presence or absence of malignancy in patient samples.

Malignant effusions are exudates containing fluid and cells in the pleural, peritoneal, and pericardial body spaces, and constitute an unequivocal sign of widespread cancer. These effusions are most commonly found in the pleural cavity with lung and breast cancers ^12^ and peritoneal cavity with ovarian cancer ^13^. Effusions offer a non-invasive sampling source that can be serially collected to capture disease evolution as it represents molecular alterations present in real-time and tissue samples may not always be available for variety of reasons. However, a significant limitation in the molecular analysis of effusions is the presence of benign background cells resulting in low tumor fractions which have been reported down to <0.1% of the total cell population ^14^. Consequently, molecular analysis such as next generation sequencing (NGS) assays are not performed in these specimens as the limit of detection is approximately 10% tumor cell fractions. Moreover, the sensitivity and specificity of cytological examination to identify malignant cells in effusions is often limited by the morphologic similarity between reactive mesothelial cells and tumor cells. Additionally, samples with low cancer cell fraction often result in low sensitivity (~60%), false negative, or inconclusive diagnoses (atypical or equivocal diagnosis) and unsuitable for downstream molecular and biomarker characterizations ^15–18^.

We applied the Deepcell platform, which uses image-based deep learning and multi-dimensional morphology analysis, to a specific proof-of-concept use case in the field of cytology, an approach we termed “SMART” (Single-cell, Multiplex, AI-based, Real-Time) cytology. We set out to increase sensitivity of carcinoma cell detection in cytology laboratory workflows by training a supervised AI model to identify and isolate carcinoma cells in malignant effusion samples. Cells classified as carcinoma were sorted by microfluidic flow and enriched cells were analyzed by common molecular characterization techniques to verify AI-enabled enrichment. Whole genome sequencing and targeted mutation sequencing of pre- and post-sorted patient samples demonstrated carcinoma cell enrichment resulted in higher sensitivity, enabling more specific clinical insight and therapeutic options.

In this study, we utilize deep learning morphological interpretations to identify and sort carcinoma cells from lung, breast, gastro-intestinal (GI) tract, and gynecological cancers to uncover actionable molecular insights in an unlabeled and unperturbed manner. Integrating clinical expertise and deep neural network models based on quantitative imaging has the potential to ultimately save lives by detecting carcinoma cells cytologists miss, isolating low-abundance cells for molecular workup, and enabling routine monitoring of minimally invasive liquid biopsies ^19–21^. While the goal of our AI methodology is not diagnosis, our study represents a proof-of-concept for integrating machine learning and quantitative morphological profiling to facilitate molecular cytology.

## RESULTS

### AI Model Training and Validation

To begin, we used a supervised deep learning approach and trained a model, termed the ‘malignant effusion AI model’, to identify carcinoma cells in a background of cells typically found in effusions such as leukocytes, histiocytes, mesothelial cells, and other immune cells. For both model training and validation, pure cell populations from benign and malignant body fluids originating from the pleural and peritoneal cavities were immunostained with a panel of antibodies to identify and distinguish cell types present in the fluids. Fluorescence-activated cell sorting (FACS) was used to isolate breast carcinoma (EpCAM+/Claudin-4+/CD45-; herein Claudin4), leukocyte (CD45+), mesothelial (CD45-/EpCAM-/Claudin4-/Calretinin+), histiocyte (CD45+/CD68+), and a collection of biomarker-negative cell types “other” (CD45-/EpCAM-/Claudin4-) to obtain pure cell populations (**Fig. 1A**). The model training set included FACS-sorted cell populations from breast cancer samples and cell lines (SK-BR-3) imaged on the Deepcell platform, yielding 2,597,910 high-resolution brightfield images of cells (**Fig. 1B)**. The model training set of images were sent to the deep convolutional neural network for multi-dimensional (>1,000 dimensions) quantitative morphological image analysis and model training, as described previously^22^. A model validation set of images, separate from images used for model training set, was created using 2,031,024 images of both cell lines (MeT-5A) and FACS purified patient-derived cell types, as described above and in Methods (**Fig. 1C**).

**Figure 1:**
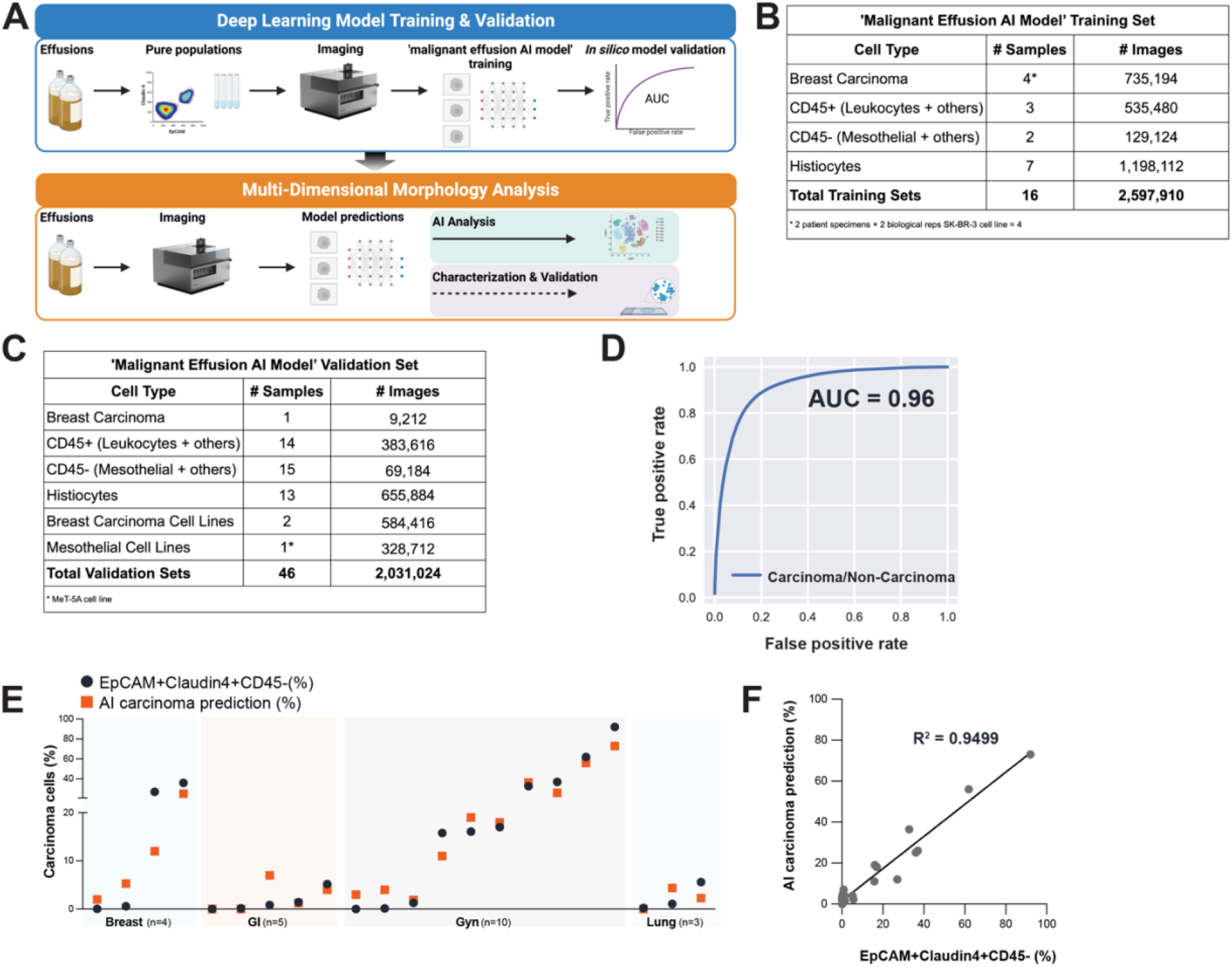
Development and Verification of AI Model for Carcinoma Cells in Effusion Samples. **(A)** Graphical schematic of experimental workflow. Top: Patient samples are labeled with biomarkers to identify and isolate breast carcinoma (EpCAM+/Claudin4+/CD45-), leukocyte (CD45+), mesothelial (CD45-/EpCAM-/Claudin4-/Calretinin+), histiocyte (CD45+/CD68+), and a collection of biomarker-negative cell types “other” (CD45-/EpCAM-/Claudin4-) to obtain corresponding pure populations using FACS. Resulting cell populations of known identity are imaged with the Deepcell instrument, and brightfield images designated as “ground truth” are input into the deep learning neural network to train the “malignant effusion AI model”. Bottom: After model training and validation, patient samples (without labels or stains) are imaged and sorted in real-time using the Deepcell instrument based on model classification. AI model predictions of carcinoma cells are then confirmed with cytology and molecular analysis. **(B)** Summary table of training sets, including clinical specimens, cells lines, and some technical replicates and number of images used for model training. **(C)** Summary table of validation sets, including clinical specimens, cells lines, and some technical replicates and number of images used for used for *in silico* model validation. **(D)** Receiver operating characteristic (ROC) curve plotting true positive rate (TPR) against false positive rate (FPR), illustrating performance of the classification model. The probability of the classification model to detect breast carcinoma cells from non-carcinoma cells is represented by the area under the curve (AUC) of 0.96. **(E)** Proportion in percent of EpCAM+Claudin4+CD45-carcinoma cells quantified by flow cytometry (gray) compared to the proportion in percent of AI model prediction of carcinoma cell (orange) in patient-derived breast, GI tract, gynecological, and lung cancer effusion samples. **(F)** Correlation plot showing proportion in percent of EpCAM+Claudin4+CD45-carcinoma cells (quantified by flow cytometry) and the proportion in percent of AI model prediction of carcinoma (quantified by AI) in effusion samples, related to **Fig. 1E**. Correlation coefficient of determination (R-squared) is 0.949.

Next, we evaluated the malignant effusion AI model performance in identifying and distinguishing breast carcinoma cells from non-carcinoma background cells with *in silico* (i.e., through computational simulation) model validation. The receiver operating characteristic curve (ROC) shows high probability (AUC=0.96) of the model identifying breast carcinoma cells from non-carcinoma cells (**Fig. 1D**). To generate a stringent ROC curve, model validation was done using all images from the validation set and each cell class was weighed such that respective contributions to the area under the curve (AUC) is the same.

Quantitative morphological features extracted from cell images are summarized as “embeddings”, which are low dimensional representations of images in vector space ^23^. These embedding vectors are not generally interpretable in terms of conventional morphology metrics but can be used to perform cluster analysis to group morphologically similar cells and visualized with Uniform Manifold Approximation and Projection (UMAP)^24^. To further validate the AI model, we generated a UMAP plot using an *in silico* mixture of known cell types including cell lines and isolated carcinoma, leukocyte, mesothelial, and histocyte cells from the model validation set (**Fig. 1C**). UMAP coloring is based on known cell identities (“ground truth”), while clustering patterns are based on AI embeddings. Breast carcinoma cells are a morphologically distinct class, which is demonstrated by separation from other cell types and clustering of carcinoma cells together (**Supplementary Fig. 1A**). Mesothelial and carcinoma cells cluster closely together with some overlap to each other, suggesting some shared morphological features between these two cell types. Histiocytes, leukocytes, and biomarker-negative “other” cells show broader distribution and higher morphological variation to carcinoma cells than mesothelial cells, as demonstrated by clear separation of these cell classes from carcinoma cells in embedding space. In addition, carcinoma cells are morphologically distinct from leukocytes which are at opposite regions of the UMAP. These common and distinct visual features detected by the AI model are consistent with morphological observations seen in effusion cell types by cytologists ^25^. Together, this UMAP of isolated known cell types comprised of 15,000 images (randomly samples from >2 million validation cell images) verify morphological features detected by the deep learning model are consistent with ground truth.

To further evaluate model performance against ground truth, we compared AI-predicted and biomarker-based (flow cytometry) carcinoma cell percentages on a cohort of pleural and ascitic effusion samples from patients diagnosed with different cancer indications. Cytologist-confirmed malignant effusion samples from patients diagnosed with breast (n=3), GI tract (n=5), gynecological (n=10), and lung (n=3) cancers were divided and processed in parallel for (1) imaging on the Deepcell platform and (2) EpCAM+/Claudin4+/CD45-biomarker immunolabeling by flow cytometry that served as ground truth. The AI model prediction for carcinoma cell percentage was highly concordant (R^2^=0.949) to flow cytometry quantification (EpCAM+/Claudin4+/CD45-), with correlation remaining linear over a wide dynamic range (0.0%-91.2%) (**Fig. 1E-F**). This indicates that although label-free, the malignant effusion AI model can detect carcinoma cells and with similar accuracy as protein expression-based biomarkers. These results further suggest that although the model was trained on breast carcinoma cells, the model learned distinct morphological features linked to carcinoma cells such that carcinoma cells from other origins (e.g., GI tract, lung, and gynecological) are distinguished from benign cells. It has been reported that the frequency of metastasis in effusion samples from GI tract (~4-6%) and lung (~6%) cancers are lower than in breast (~40%) and ovarian (~22%) cancers ^26^. Our results are relatively consistent with these clinical findings, with predicted carcinoma cells frequencies in GI tract and lung cancers lower (mean, median, range) compared to breast and gynecological cancers. Importantly, the AI model was able to correctly predict the proportion of tumor cells at low (~0%) and high (~80%) ends of tumor frequency (**Fig. 1E, Supplementary Fig. 1B-C**). Therefore, the lower carcinoma percentage reported in these samples reflect true biology for these cancer types. Differences in carcinoma frequency quantification between biomarker-based flow cytometry and AI model observed were not statistically significant (**Supplementary Fig. 1B**). Together, these data suggest the AI model can detect carcinoma cells from diverse cancer types and independently of its tissue of origin over a wide dynamic range.

### Morphologically Similar Cell Types Cluster Together in AI Embedding Space

To assess the morphology of carcinoma cells between patients, we applied the trained and validated malignant effusion AI model to generate cell class predictions in specimens from two breast cancer patients. Patient A (25.2% AI carcinoma prediction) was diagnosed with Her2-metastatic breast cancer and Patient B (42% AI carcinoma prediction) was diagnosed with ER+/PR+/Her2 indeterminate invasive lobular breast cancer. AI embedding UMAP analysis shows carcinoma cells from Patient A and B cluster separately from mesothelial, leukocytes, and histiocytes, which cluster together within each respective cell class and suggest stability of embeddings across the two patients for non-carcinoma cells (**Fig. 2A**). Interestingly, carcinoma cells from these two cases cluster separately from each other, indicating morphologic dissimilarity of carcinoma cells between different patients (**Fig. 2A**). Indeed, representative brightfield images of carcinoma cells confirm differences detected by the AI model, with Patient A cells exhibiting more granularity and textured appearance compared to Patient B cells (**Fig. 2A**), reinforcing that the cells are morphologically distinct.

**Figure 2:**
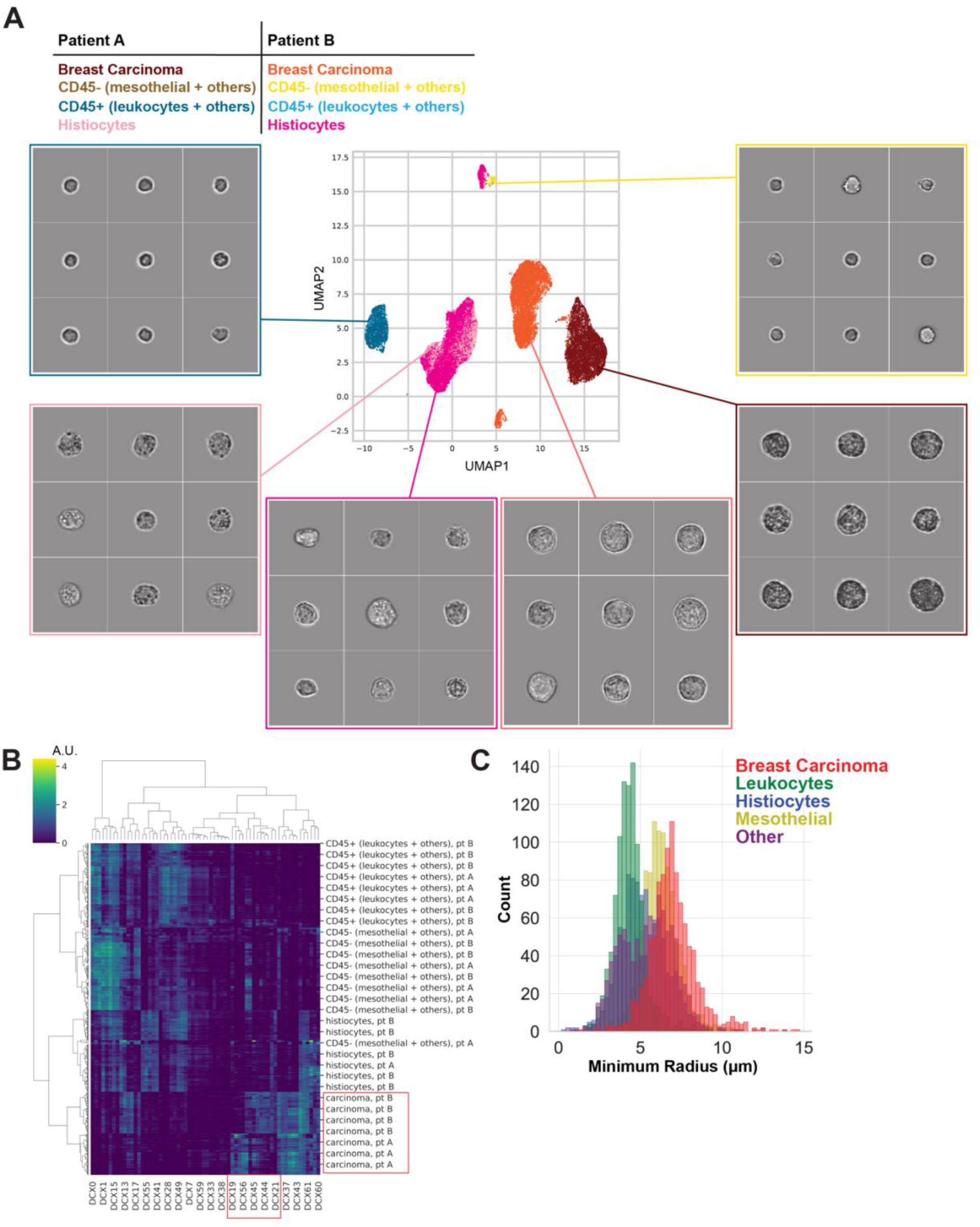
Morphologically Similar Cell Types Custer Together in AI Embedding Space. **(A)** Embeddings UMAP plot of AI prediction of cell classes. Embeddings are extracted from images of unlabeled breast cancer effusion samples from two different patients. Classifications of cell types are based on the malignant effusion AI model. Distinct cell classes are shown, including breast carcinoma (red/orange color family), CD45-(mesothelial + others) (gold color family), CD45+ (leukocytes + others) (blue color family), and histiocytes (pink color family). Representative brightfield images from each cluster are shown; for display only, these images have been normalized for illumination intensity, and the pixels outside the cells have been blanked out. **(B)** Dendrogram heatmap of embedding vectors with highest contribution to AI prediction of cell classes. Red box highlights vectors (DCX19, DCX56, DCX45, DCX44 and DCX21) contributing to distinct clustering of carcinoma cells from Patient A and B. **(C)** Histogram plot by isolated known cell types, as quantified by computer vision morphometric measures of cell size (radius). Isolated known cell types (“ground truth”) are as follows: breast carcinoma (EpCAM+/Claudin4+/CD45-), leukocyte (CD45+), mesothelial (CD45-/EpCAM-/Claudin4-/Calretinin+), histiocyte (CD45+/CD68+), and biomarker-negative cell “other” cells (CD45-/EpCAM-/Claudin4-).

To gain more insights into cell features contributing to AI cell predictions, we analyzed the embedding vectors and the respective contribution for each cell class (**Fig. 2B**). Each cell class exhibits unique profiles, with heatmap color representing embedding value. Several AI feature vectors (DCX19, DCX56, DCX45, DCX44, DCX21) differentially contributing to carcinoma predictions between Patient A and B (**Fig. 2B**). Since carcinoma cells are generally larger than background cells present in body fluids, we next asked whether cell size was the distinguishing feature used in AI cell class predictions. Computer vision measurements of minimum radius indicate overlap in size between all cell classes, with carcinoma cells having highest overlap with mesothelial cells followed by histiocytes, biomarker-marker negative “other” cells, and leukocytes (**Fig. 2C**). This indicates that the model learned morphology features other than cell size to classify carcinoma from non-carcinoma cells.

To further compare the morphology of carcinoma cells from breast cancer patients, we added a third breast cancer patient to the embeddings analysis (**Supplementary Fig. 2A**). Patient A and C carcinoma cells plot as distinct clusters but close together, suggesting high relative morphological similarity compared to Patient B carcinoma cells. Representative carcinoma cell images from Patient A show cells exhibit more granularity and texture, while cell images from Patient C show cells are relatively larger with medium granularity and texture (**Supplementary Fig. 2A**). We applied computer vision morphometric measurements to gain further insight into morphological profiles, namely spot measurements and texture characterization. The spot measurement isolates large dark sports in a specific size range using the Laplacian of Gaussian filter ^27^ and the area opening ^28^. For texture characterization, we used the uniform local binary pattern (LBP) variant ^29,30^. Computer vision morphometrics confirmed Patient A cells had more large dark spots compared to Patients B and C (**Supplementary Fig. 2B**). Additionally, morphometrics verified larger cells in Patient C and a different texture in the cell’s periphery in Patients A and C compared to Patient B (**Supplementary Fig. 2B**). These results are consistent with visual examination from representative images, and morphometric measurements hint at morphological differences underlying interpatient heterogeneity (**Supplementary Fig. 2A-B**).

### Processed Cells are Viable and Compatible with Cytology Workflows

One of the major challenges of using effusion samples as liquid biopsy material is retrieving viable and intact cells for molecular and functional analyses. Many enrichment methods rely on fixation, epitope tags, or other techniques that may alter and destroy cells during the process. We previously reported cell lines and dissociated tumor tissue run through the Deepcell platform retain viability and gene expression profiles^22^, but have not tested clinical effusion samples. We evaluated cell viability of breast and ovarian cancer pleural fluid samples by trypan blue exclusion with and without instrument processing. Paired t-testing demonstrated no significant differences in viability after instrument processing, which is consistent with our previous results^22^ (**Fig. 3A**).

**Figure 3:**
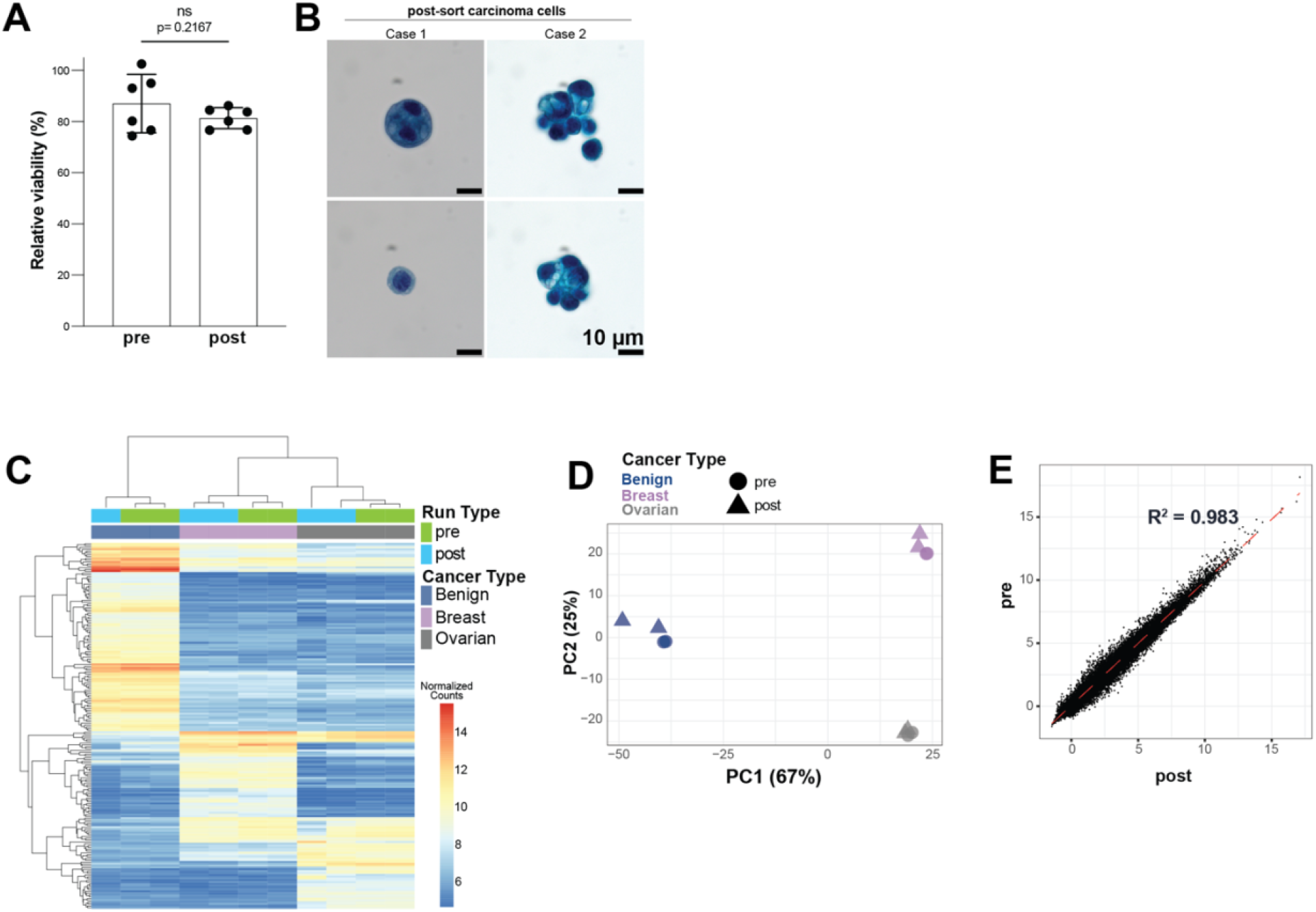
Processed Cells are Viable and Compatible with Cytology Workflows. **(A)** Relative cell viability of samples before (pre) and after (post) instrument flow-through. **(B)** PAP stain performed and assessed by a board-certified pathologist on two high-grade serous ovarian cancer ascitic effusion samples show sorted cells remain intact and amenable to cytology staining. Scale bar as indicated. **(C)** Bulk RNA-Seq heatmap showing normalized gene expression counts of pre (green) and post (blue) instrument flow-through effusion samples from patients diagnosed with non-cancerous disease (benign; dark blue), breast (purple), and ovarian (grey) cancers. **(D)** Principal component analysis (PCA) plot of pre (circle) and post (triangle) instrument flow-through non-cancerous disease (dark blue), breast (purple), and ovarian (grey) cancer samples. Principal component 1 (PC1) represents 67% contribution of variation, and principal component 2 (PC2) represents 25% contribution of variation. Related to **Fig. 3C**. **(E)** Correlation plot showing pre versus post instrument flow-through sample gene expression based on bulk RNA-Seq analysis. Each dot represents the transcript count from pleural fluid derived from patient diagnosed with ovarian cancer fluid (X- and Y-axis is transformed (regularized logarithm; rlog) count from pre and post flow-through samples, respectively). R square value is indicated in the graph. Related to **Fig. 3C**.

To test whether sorted cells are intact and retain morphological architecture that cytologists use to access cell identity, ascitic fluids were collected from two different cases of high-grade serous ovarian cancer for Papanicolaou (PAP) staining after sorting with 0.85 AI confidence threshold. Samples were processed with cytospin, mounted, and stained on slides (**Fig. 3B, Supplementary Fig. 3**) then examined under a microscope by a board-certified pathologist. Results demonstrated that cells retain morphological architecture and are compatible with cytological staining, indicating feasibility for applying this framework in clinical workflows.

To assess whether cells loaded and flown-through the instruments are not adversely affected and maintain gene expression profiles, we performed bulk RNA-Seq on non-cancerous disease (benign), breast, and ovarian cancer pleural effusions (**Fig. 3C-D**). For each cancer type, gene expression analysis shows minimal differences between the initial samples and after instrument processing, suggesting cells run through the instrument are not affected or altered (**Fig. 3C**). The major variability observed in gene expression is driven by sample-specific signatures rather than instrument processing and sample handling (**Fig. 3D**). Gene expression correlation coefficient (R^2^) between gene expression profiles pre- and post- flow-through was 0.983, signifying that cells processed through the Deepcell instrument for downstream analysis are directly comparable with unprocessed cells (**Fig. 3E**).

### Validation of AI-mediated Identification of Carcinoma Cells Using Molecular Analysis

To confirm this framework is feasible for cytology workflows, pancreatic adenocarcinoma, high-grade serous endometrial carcinoma, and high-grade ovarian/fallopian tube carcinoma ascitic fluids were collected for PAP staining before and after cell sorting (**Fig. 4A, Supplementary Fig. A-B**). Pre- and post-sort samples were sent to a board-certified pathologist for analysis. Results confirmed a higher purity of cells of interest and atypical cells, with depleted inflammatory cells eliminating visual noise on the slides. This confirms AI carcinoma predictions and platform sorting enables higher sensitivity for cytopathology analysis.

**Figure 4:**
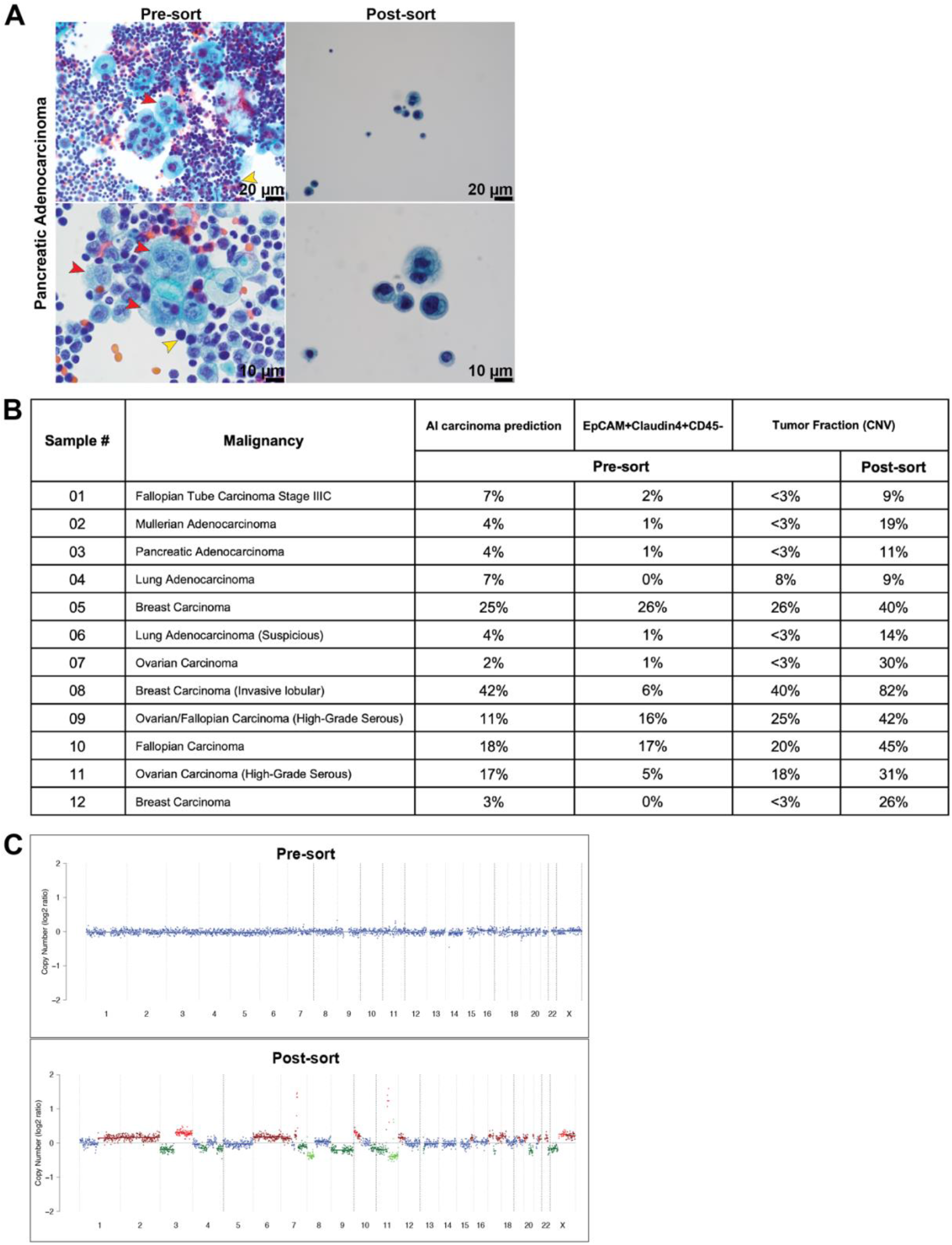
Validation of AI-Driven Identification of Carcinoma Cells. **(A)** PAP stain performed and assessed by a board-certified pathologist on pre- and post-sort pancreatic adenocarcinoma ascitic fluid at different magnifications. Red arrows indicate cells of interest/atypical cells, yellow arrow indicate inflammatory cells. Scale bar as indicated. **(B)** Table of sample cases indicating pre-sort proportion of carcinoma cell in percent predicted by the AI model (AI carcinoma prediction), proportion of carcinoma cell (EpCAM+/Claudin4+/CD45-) measured by flow cytometry, and tumor fractions calculated from CNV analysis. Corresponding post-sort tumor fractions are shown for samples after AI carcinoma cell enrichment. **(C)** Representative CNV plots showing amplifications/deletions in pre- and post-sort patient samples.

Thus far, we have shown (1) AI carcinoma cell predictions for all cancer types are highly concordant to protein expression-based identification of carcinoma cells by flow cytometry, (2) AI image analysis shows that carcinoma cells are morphologically distinct cells, and (3) carcinoma cells cluster separately from non-carcinoma cells in embedding space. To more definitively verify that sorted AI predicted carcinoma cells are malignant, we supplemented our analysis with molecular methods to assess and characterize cells. We performed whole genome sequencing (WGS) followed by copy number variation (CNV) analysis on patient samples with low (0%) to mid (25-42%) carcinoma frequencies to cover a range of initial pre-sort effusion compositions (**Fig. 4B**). Results show high concordance between AI carcinoma predictions and %EpCAM+Claudin4+CD45-, reaffirming consistency of the AI model with protein expression (ground truth). Further, pre-sort samples exhibit high concordance with genomic assessment of tumor fraction based on CNV analysis (**Fig. 4B**). After sorting, all samples demonstrated an increase in tumor fraction after carcinoma cell enrichment, as analyzed by ichorCNA (**Supplementary Fig. 4C)**^31^. One exception is Sample #4, which had a primary cytology diagnosis of <1% malignant and 0% by flow cytometry, suggesting the initial processed sample may not have contained malignant cells. Of note are Samples #8 and #11, which were diagnosed with invasive lobular breast carcinoma and high-grade serous ovarian carcinoma, respectively. Sample #13 carcinoma frequency was estimated at 42% by AI and 6% by flow cytometry (EpCAM+Claudin4+CD45-), but the estimated tumor fraction at 40% was more concordant with AI. Sample #16 was estimated at 17% by AI, 5% by flow cytometry, and 18% by tumor fraction. These data suggest deep morphological interpretation may overcome label and epitope-based constraints such as biomarker specificity and provide a more comprehensive screening for malignancy. Remarkably, all cases demonstrated enriched CNV profiles and at least 2X increase in tumor fraction with undetectable^31^ (<3%) tumor fractions and post-sort increases to detectable levels (**Fig. 4B-C**). Together, this indicates AI carcinoma predictions are concordant with protein and genomic expression and accurate in carcinoma cell classification and sorting, as validated by WGS. Consequently, this augmented workflow enabled detection of actionable molecular information that was not detectable in the original patient sample.

### Targeted Mutation Analysis Reveals Increase Sensitivity for VAF Detection

Lastly, targeted mutation sequencing analysis was performed on select patient samples to verify enriched populations yielded higher sensitivity for mutational detection. Sorted carcinoma cell samples showed striking improvement in somatic variant allele frequency (VAF) in six of the seven patient cases (**Fig. 5**). The seven patient cases were selected based on the availability of primary tumor mutation profiles of donor samples processed on the Deepcell platform. The primary mutation profile panel was obtained independently using a 50-gene cancer hotspot panel. Using a different 56 oncogene panel, we established the somatic VAF before and after carcinoma cell sorting of the effusion samples on the Deepcell platform. Results showed mutations detected in post-sort samples match mutations detected in the primary tumor tissue and/or liquid biopsy specimen, affirming that the platform accurately identifies and enriches for carcinoma cells with clinically relevant somatic mutations from diverse cancer types.

**Figure 5:**
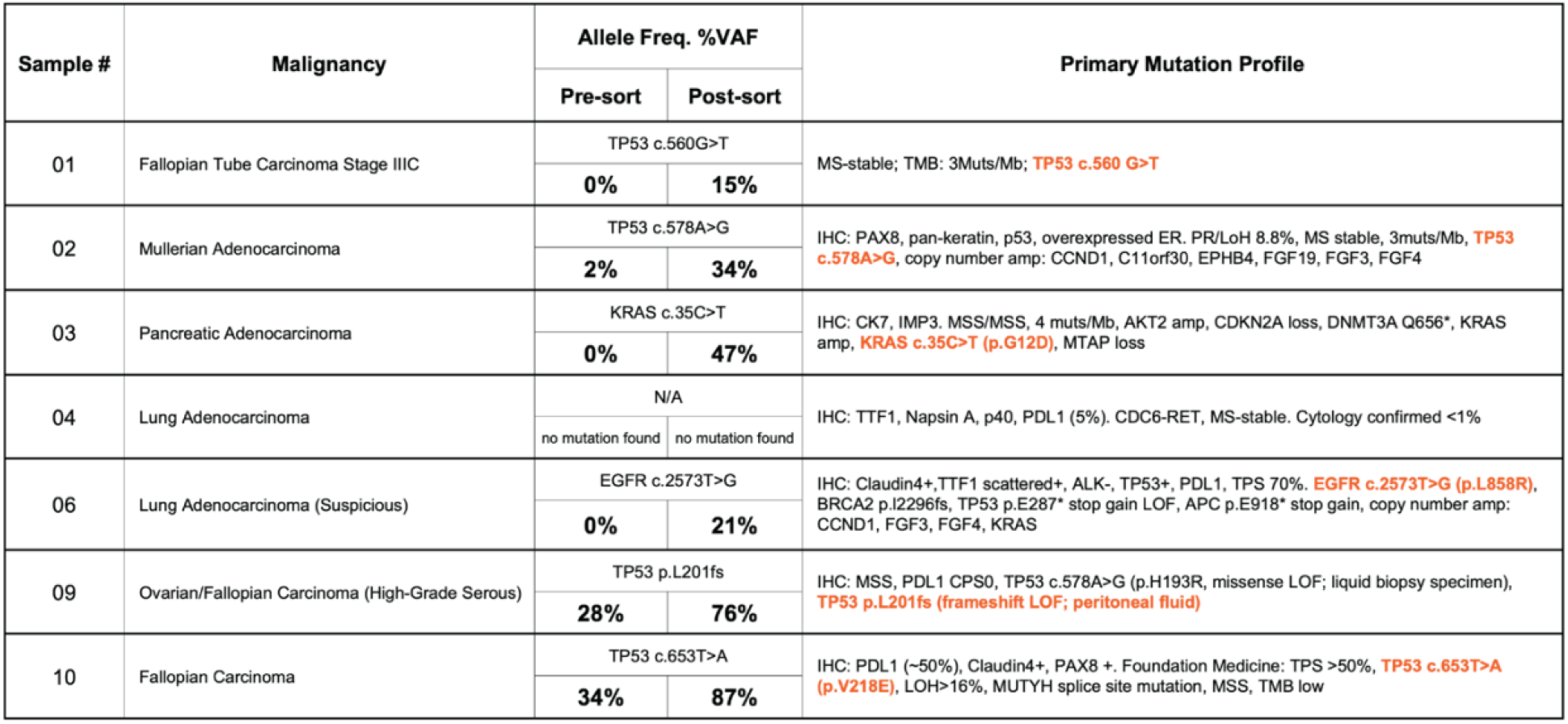
Targeted Mutation Analysis Reveals Increased Sensitivity for VAF Detection. Table of sample cases showing pre- and post-sort somatic variant allele frequency (VAF), as detected by targeted mutation sequencing (56G oncogene panel). Primary mutation profiles from tissue and/or IHC for corresponding sample cases are shown, with orange highlighted text indicating matching detected mutation in post-sort specimens determined by Ampliseq Cancer Hotspot panel V2.

Of note, mutations that were undetectable (<1%) in three pre-sort specimens were identified at VAFs ranging from 15% to 47% after sorting. Two pre-sort specimens with mutations at VAFs lower than the typical limit of detection for conventional next generation sequencing demonstrated post-sort VAFs of 15% and 34%. For example, EGFR c.2573T>G (p.L858R), a known pathogenic somatic mutation in lung adenocarcinoma, was detected at 21% VAF in the post-sort sample but not in the pre-sort patient sample (**Fig. 5**). These cases with pathogenic VAF detection in post-sort but not pre-sort samples correlating with primary tumor tissue further supports integration of AI enriched effusion samples with cell cytology workflows to facilitate more sensitive and efficient molecular analysis. Thus, the application of deep learning to morphological characterization of malignant effusions has the potential to enable insight into actionable molecular lesions and inform treatment options.

## DISCUSSION

Cytology is a well-established first line of diagnostic and screening method for human diseases, especially cancer. The field underwent several generations of development, from traditional smear-based cytology to liquid-based (monolayer preparation) cytology to currently digital and computational cytology. However, even with advances of digital and molecular technologies, molecular-driving cytological analysis is still limited due to the lack of robust, standardized techniques for preparing cytology samples for molecular analysis. Additionally, current morphology-based cytological analysis is qualitative, subjective, and restricted by features the human eye can define, with a mean sensitivity of 60% depending on the primary tumor, sample preparation, and individual cytologist ^32^. Furthermore, molecular analysis of carcinoma cells is needed to identify actionable targets ^33–36^. Morphology-based cytology analysis alone, even with the addition of ancillary techniques such as immunohistochemistry or cytogenetic analysis ^37–39^, is insufficient for achieving this goal given the increasing number of targets in the context of tumor heterogeneity ^40–43^.

Methodologies currently exist to increase tumor cellularity before molecular testing is performed. With liquid-based cytology where effusions are directly transferred to glass slides or used to create cells blocks from which microscopic sections are subsequently cut and transferred to glass slides, select areas with focally increased tumor content can be manually scrapped off and submitted for downstream processing ^44^. Laser capture microdissection is a more sophisticated method involving the use of a laser to dissect out small regions or even single cells more precisely. As these methods depend on morphologic assessment by a human operator, similar issues exist as with cytological evaluation, and they are inefficient when tumor cells are present in low quantities or difficult to distinguish from benign background cells. For instance, it is historically difficult to distinguish mesothelial cells from carcinoma cells in effusion samples based on morphology, immunohistochemistry (IHC), and/or cytological stains. Though there may be some subtle features (e.g., mesothelial cells often show “windows” and “hugging” between cells, etc.), an individual mesothelial cell, especially when it is reactive (e.g., stimulated by inflammation) can be difficult to distinguish from a metastatic carcinoma cell, which may result in an equivocal or atypical diagnosis ^45^. As carcinoma cells may be limited in number, even using a battery of IHC markers may not provide a definitive answer, posing a diagnostic challenge. Accordingly, applying machine intelligence as a facilitating tool can aid in detecting visually problematic cells such as mesothelial versus carcinoma cells in a robust manner, as demonstrated in this study. Of note, we were able to enrich for unlabeled carcinoma cells to the extent that molecular changes below the standard limit of detection of current next generation sequencing (approximately 5%) were detected at high levels ranging from 14% to 47% allele frequencies. These results highlight the potential of the methodology to isolate rare single cancer cells from a background of benign cells with applications in minimal residual disease detection ^46,47^.

One limitation of this study is variable levels of enrichment seen in several cases. We posit this issue can be attributed to two factors: (1) physical hardware, such as the performance and speed of pneumatic sorting valves and (2) model performance, such as whether enough heterogeneity of cell types and effusion specimens were used to train the model. In future iterations, mechanical improvements and adjustments in microfluidic flow rates can be made to increase purity levels. Furthermore, additional biological diversity of patient specimens from different cancer types and varying carcinoma frequencies may enhance model performance and enrichment levels. For instance, in this study we observed partial overlapping of carcinoma and mesothelial cells as observed in the AI embeddings UMAP clustering, which may result in decreased depletion of mesothelial cells compared to other non-carcinoma cells. To reduce the degree of overlapping, we expect that the model can be trained further with additional annotated cell images of cell line and patient-derived mesothelial cells to discern carcinoma and mesothelial morphological differences. We interpret this limitation as related to the same cytopathologist challenge in distinguishing morphologically similar cells such as carcinoma cells and mesothelial cell compared to smaller lymphocytes, even with aid of immunohistochemical stains.

Applying advancements in artificial intelligence (AI), specifically deep learning, to tissue-based images has become an intense area of research in digital pathology with applications including cancer detection ^48,49^ and staging based on the tumor architecture ^50,51^, among others. Further support for this paradigm shift in precision medicine was demonstrated in a recent study showing the combination of AI and clinician detects more instances of breast cancer than either alone ^19^. However, AI has not been extrapolated to single-cell cytology analysis such as label-free tumor cell enrichment. Here we present an AI-driven technology platform to classify and isolate carcinoma cells from breast, lung, GI tract, and gynecological cancer effusions using learned morphological representations. Interestingly, although the AI model was trained on breast carcinoma cells (cell line and isolated patient cells), it performed remarkably well on other cancer types assessed in this study (lung, GI tract, gynecological). This indicates carcinoma cells residing in effusions share similar morphological properties, which may potentially be associated with metastatic fitness and capacityl^6,7^.

Importantly, as proof-of-concept for precision medicine applications, while critical molecular events such as clinically relevant actionable mutational targets were undetectable without processing, such mutations were detected after processing. Microfluidic sorting, although not currently highly efficient, allows effusions to retain viability and compatibility with genomic analysis such as RNA-seq and WGS, making sorted cells suitable for a comprehensive assessment of patient’s mutation profile. We additionally validated sorted cells are compatible with downstream single-cell RNA-Seq and live cell culture for drug testing and other experimental conditions (data not shown). Beyond the physical enrichment of target cells, high-content images are captured and stored, allowing for continuous measurement and re-analysis of captured images to detect additional phenotypes resulting from differing cell states/types, drug treatments, or molecular aberrations at single-cell resolution. This technology provides the foundation for the future practice of SMART cytology.

## MATERIALS AND METHODS

### Specimen Collection

Ascites and pleural effusion specimens were collected from patients admitted to UCLA Medical Center (Los Angeles, CA) and reviewed by trained pathologists or cytologists. As byproducts of diagnostic procedures, samples were de-identified and the study was exempted from the UCLA Institutional Review Board. Specimens with >100mL leftover fluid following cytological diagnostic analysis were shipped to Deepcell (Menlo Park, CA). Samples were stored at 4°C within 24 hours of collection, shipped on ice, and processed within <48 hours from time of collection. Specimen information including time of collection, treatment history, date of treatment, cytopathology and molecular diagnostic results, histological diagnosis, and percentage of malignant cells determined by cytology was collected and shared. Additional patient specimens were commercially obtained from Bio-Options (Brea, CA, USA) and Discovery Life Sciences (DLS; Los Osos, CA, USA).

### Sample Preparation

Upon receipt, effusion samples were centrifuged to pellet the cells. The cell pellet was resuspended in Ham’s F-12K (Kaighn’s) Medium (Gibco) + 10% Fetal Bovine Serum (Peak Serum) and strained using a 40μm cell strainer (Corning), then inspected for cell clumps. If clumps were present in the strainer, 1mg/mL collagenase was used for dissociation (Sigma-Aldrich). Combined cells and dissociated clumps were then fixed with 4% paraformaldehyde (EMS) for 20 minutes at room temperature with gentle agitation and resuspended with calcium- and magnesium-free PBS (Gibco). If RBC were present, the cell suspension was stored overnight at 4°C to allow spontaneous hemolysis then washed with PBS. Fixed samples were stored at 4°C until use. Single cell suspensions and removal of red blood cells (RBC) was performed with RBC lysis buffer diluted in water according to manufacturer’s recommendations (BioLegend). For cryopreservation, when applicable, all steps above were performed, cells were then washed and resuspended in CryoStor® CS10 (Stemcell Technologies). For cell viability assessment, pre-sort or post-sort cells were stained with trypan blue (Gibco). Cells were then counted under a microscope (Leica).

### Cytology

We used conventional cytology methods to perform cytomorphological analysis of body fluids samples. Briefly, an aliquot (50 mL) of effusion fluid sample was transferred to a 50 mL conical tube and centrifuged. The supernatant was decanted by manual pipetting, so as to retain 3-4 mL of pellet mixing well with supernatant remaining in the tube to make cytospin slides and cell block. For samples containing red blood cells indicated by pink to red colored specimens, the cell pellet was overlaid with 10 mL of CytoRich Red (BD), resuspended, and centrifuged again. After assembling Shandon cytospin slide into the cytospin processing bin, one to three drops of cell pellet suspension were added into each cytospin chamber. After securely fasten the lid onto the processing bin (“Seated head roter”), the cystospin bin was placed in the Cytospin processor (Richard-Allan Scientific) and centrifuged at 191 g for 5 minutes. Slides were immediately fixed in 95% ethanol for 20 min. The staining process was carried out using a Tissue-Tek Prisma® Plus Automated Slide Stainer (Prisma-P-AD, Sakura Finetek USA). For cell block preparation, cells were fixed by adding 1 mL of zinc formalin to the remaining cell pellet, mixed by pipetting, and centrifuged. After stood for 1 hour, the material was transferred into a biopsy bag, and submitted to the histological laboratory for paraffin embedding, sectioning (4 to 6 micron), and routine Hematoxylin and Eosin (H&E) staining. All prepared slides were reviewed and examined by clinical technologists and board-certified pathologists under the light microscope (BX53, Olympus).

### Cell Viability

For cell viability assessment, pre-run (T = 0 hr), post-run (flown-through + T = 3 hr), and “un-run” (room temperature for T = 3 hr) cells were stained with trypan blue (Gibco). Cells were then counted under a microscope (Leica). Relative cell viability was calculated between post-run/pre-run cells and un-run/pre-run cell viability.

### Sample Preparation for Model Training and Validation

For AI model training and validation, pure populations of known cells were be obtained by FACS isolation of CD45+ (leukocyte, Biolegend Cat#304006), CD45+/CD68+ (histiocytes, BioLegend, Cat#333827), Calretinin (mesothelial, Santa Cruz Biotechnology Cat#sc-365956), and EpCAM+/Claudin4+ (carcinoma cells, R&D Systems Cat#AAGV0320031). Fixed cells isolated from patient samples were stained with respective primary antibodies for 30 minutes at 25°C and washed twice with BD Stain buffer (Becton-Dickinson, Cat#554656). Cells were FACS sorted on a BD Melody (Becton-Dickinson) using the following gating strategy: First, debris is eliminated by FSC-A vs. SSC-A, then cell clumps and doublets are removed by SSC-H vs. SSC-W and FSC-H vs. FSC-W. Leukocytes are identified by CD45 expression and gated further by CD45+ vs. CD68 with histiocytes identified by CD68 expression. The CD45-population is investigated by Claudin4 vs. EpCAM for cancer which is determined by both Claudin4 and EpCAM expression and the population with no expression from Claudin4 and EpCAM being defined as CD45-other population which includes other non-hematopoietic and epithelial cells. The CD45-other population is then gated by CD45-vs. Calretinin with mesothelial cells identified by Calretinin expression. In addition to patient samples, commercially available cell lines SK-BR-3 (ATCC, Catt# HTB-30™) and MeT-5A (ATCC, Cat# CRL-9444™) maintained per manufacturer’s recommendation were used for model training and validation.

### Model Training and Validation

#### Instrument

The Deepcell instrument consists of hardware (microfluidic chip, high-resolution imager, real-time signal processing), deep learning convolutional neural network model^52^, and proprietary software. Captured images are cropped around the cell and fed into a convolutional neural network for generation of multi-dimensional morphological descriptors and classification in real-time. Based on cell classification, pneumatic sorting valves are activated for either target cell collection or waste. Target cells are classified using a confidence threshold level over 0.85 (based on the softmax activation function^53^, which transforms the raw neural network outputs into probability vectors) and collected. Cells below the set confidence threshold and other cell classes go to waste. Sorted cells are then retrieved for downstream analysis.

#### Dataset

To develop the ‘malignant effusion AI model’, pure cell populations (cell lines, FACS-sorted patient samples) were divided into multiple aliquots and imaged on more than one instrument to mitigate inter-instrument variations on the model training. Pure cell populations were used to curate training and validation sets composed of the following cell classes: breast carcinoma, CD45+, CD45- and histiocytes. Distinct samples and cell lines were used for the training and validation sets. For each cell class, we manually annotated “out of focus” images (cell that is blurred and intracellular features are not sharp or clearly defined) and debris/non-cellular images (bubbles, blank frames, unhealthy cells, cell clumps) to exclude images that may cause misclassification.

#### Classification model

All cell images were resized to 299×299×3 pixels to make them compatible with the InceptionV3 architecture^52^. The training progressed 30k steps with a batch size of 1024, and a 0.16 learning rate on an Adam optimizer. The training algorithm was modified to consider the skewed distribution of cell classes by providing class weights which are inversely proportional to the class frequencies in the training data. The classifier thus learns equally from all classes.

#### Embedding model

We implemented a variant of the InceptionV3 model to construct the embedding model. From one of the top layers proximal to the softmax output layer, we took one of the final dense layers and input it into a size 64 embedding layer. This embedding InceptionV3 model was trained with the same hyperparameters as in the supervised classification model training.

### Bulk RNA-Seq

Total RNA was extracted from cells using the RNeasy Mini Kit (Qiagen). cDNA synthesis, amplification, and library preparation were performed with the Quantseq 3’m RNA-seq Library Prep Kit (Lexogen) according to the manufacturer’s protocol. The final libraries were sequenced on the Illumina Miniseq system using 150-cycle MiniSeq High Output Reagent Kit (FC-420-1002). The demultiplexed reads in FASTQ format were trimmed for adapter sequence using the Atropos^54^ software with Illumina universal primer read sequence and the trim parameters, where the maximum trimmed length is set at 50bp and quality score cutoff is set at 20, with the option of discarding reads containing more than 5 N bases and removing flanking N bases from each read ^54^. The trimmed reads are then aligned to the human genome, GRCh38.p13 using STAR aligner ^55^. The raw count for each gene is generated, where the Ensembl (https://ensembl.org; European Bioinformatics Institute) gene ID is used as an identifier. For downstream gene expression analysis, the raw count data from each individual sample is aggregated into a single raw count table. Differential expression analysis was done using DESeq2 ^56^.

### WGA and CNV Preparation

Whole genome amplification (WGA) was performed using ResolveDNA™ WGA (BioSkryb), according to manufacturer’s protocol, on 500-1000 both presorted and Deepcell-sorted cells. NGS libraries were constructed using WM kit (Watchmaker Genomics), 500 ng of WGA product were used to prepare a library for sequencing with Watchmaker DNA Library Prep Kit with Fragmentation. The library was sequenced for copy number variation (CNV) analysis on Illumina NextSeq 1000 (Illumina) with NextSeq1000/2000 P2 Reagents (300 Cycles) v3 reagent (Illumina).

### CNV Analysis Method

The raw FASTQ files are aligned to the human reference genome, GRCh38.p13, using BWA-MEM algorithm by bwa software, version 0.7.17 ^57^. The aligned reads are checked for duplication, and those reads with the alignment quality score 20 and above are selected for the CNV analysis. The read count coverage files for each sample are generated in WIG format using HMMCopy’s readCounter command with 1Mb bin size option. The panels of normals are prepared from the benign cell lines, GM12878, GM24143, GM24149, GM24385, and HFL1, to be used as a reference during the CNV analysis, following the recommended processing steps provided by ichorCNA manual. These files are used as an input to ichorCNA software (bioconda r-ichorcna version 0.3.2), for CNV analysis and tumor fraction prediction ^31^, using the ranges of ploidies of 1.75, 2, and 2.25 and initial normal contamination percentage ranging from 1% to 99%. The solutions ranked by the descending log-likelihood values by the ichorCNA are manually reviewed to eliminate the suboptimal solutions and to identify the solution that best explains the data, as recommended by ichorCNA.

### Spike-in Data CNV Analysis

To evaluate the tumor fraction prediction performance from ichorCNA on the whole genome sequencing data generated and processed at Deepcell, a benign cell line, GM12878 and lung adenocarcinoma cell line, NCI-H23, were mixed at various ratio with expected tumor fractions ranging at 5, 10, 20, 50% using 25, 50, and 100 cells. Whole genome amplification was performed on the prepared mixture samples with BioSkryb ResolveDNA™ Whole Genome Amplification Kit (BioSkryb Genomics) and Takara PicoPlex WGA v3 kit (Takara) following the manufacture’s protocols. The libraries were prepared using Roche KAPA HyperPlus Kit (Roche) with 500 ng WGA product input and sequenced with iIlumina Nextseq 1000 (Illumina). The observed tumor fraction and expected tumor fraction were compared and fitted using the linear regression model.

### Mutation Analysis with Ion AmpliSeq Cancer Hotspot Panel v2

Targeted amplification of multiple coding regions of 50 genes recurrently mutated in solid tumors (*ABL1, AKT1, ALK, APC, ATM, BRAF, CDH1, CDKN2A, CSF1R, CTNNB1, EGFR, ERBB2, ERBB4, EZH2, FBXW7, FGFR1, FGFR2, FGFR3, FLT3, GNA11, GNAQ, GNAS, HNF1A, HRAS, IDH1, IDH2, JAK2, JAK3, KDR, KIT, KRAS, MET, MLH1, MPL, NOTCH1, NPM1, NRAS, PDGFRA, PIK3CA, PTEN, PTPN11, RB1, RET, SMAD4, SMARCB1, SMO, SRC, STK11, TP53*, and *VHL*) was performed using the Ion AmpliSeq Cancer Hotspot Panel v2 (ThermoFisher Scientific, Waltham, MA). Following next-generation sequencing on the Ion Torrent PGM system (ThermoFisher), the raw sequencing data was processed using the Torrent Suite Software and Ion Reporter Software with resultant bases aligned to targeted regions (hg19). Variant call format (VCF) files were generated with the variantCaller plugin. Detected alterations were filtered and reviewed manually to determine pathogenicity and reportability. Depending upon the total amount of DNA present, the assay can detect ~5% of an alternate allele in a background of reference alleles. This method is limited to the detection of certain categories of mutations, including single nucleotide variants and small insertions/deletions (≤18 nucleotides), within the regions of the genome targeted by this assay.

### Mutation Analysis with xGen™ 56G Onco Amplicon Panel v2

Target amplification for mutation detection was performed using 20 ng of WGA product with Integrated DNA Technologies xGen™ Amplicon Core kit and xGen™ 56G Onco Amplicon Panel v2 (Integrated DNA Technologies), which offers comprehensive and hotspot coverage of 56 clinically-relevant oncology-related genes (ABL1, AKT1, ALK, APC, ATM, BRAF, CDH1, CDKN2A, CSF1R, CTNNB1, DDR2, DNMT3A, EGFR, ERBB2, ERBB4, EZH2, FBXW7, FGFR1, FGFR2, FGFR3, FLT3, FOXL2, GNA11, GNAQ, GNAS, HNF1A, HRAS, IDH1, IDH2, JAK2, JAK3, KDR, KIT, KRAS, MAP2K1, MET, MLH1, MPL, MSH6, NOTCH1, NPM1, NRAS, PDGFRA, PIK3CA, PTEN, PTPN11, RB1, RET, SMAD4, SMARCB1, SMO, SRC, STK11, TP53, TSC1, VHL). Target amplicons were sequenced using Illumina Miniseq with Miniseq Mid Output Kit (300-cycles) reagent (Illumina). Depending on DNA input and sequencing depth, the assay can detect ~1% of an alternate allele in a background of reference alleles.

### Mutation Analysis Method

The Illumina adapter-trimmed FASTQ files are aligned to the human reference genome, GRCh38.p13, using BWA-MEM algorithm, followed by the clipping of amplicon primer-binding regions by ampliconclip, template length correction by fixmates and *MD* and *NM* tag fix by calmd, all available from samtools software, version 1.14 ^58,59^. The germline calls are made using the pileup data, using the aligned reads that overlap with the target regions of the given assay. The initial calls are made against the reference genome sequences, followed by the manual review using the IGV viewer visualization tool to eliminate the false positive calls. The somatic calls are made using Mutect2 tools from GATK4 version 4.2.2.0 ^60–62^ on amplicon target sites, using the matched normal sample. When a matched normal sample is unavailable, Mutect2 is used with the panels of normals, generated from 1000 Genomes Project and gnomAD population allele frequency VCF file. All somatic calls are further filtered using GATK4 FilterMutectCalls, with minimum unique reads supporting the alternate allele set to 2, minimum alt reads required on both forward and reverse strands set at 2, and minimum allele fraction required set at 0.01, followed by the manual review using IGV visualization tool.

### Statistical Methods and Hypothesis Testing

When applicable, statistically significant differences between experimental conditions were tested with GraphPad Prism Software (Version 9.4.0).

## AUTHOR CONTRIBUTIONS

Research design and study concept: CJ, AM, JR

Experiments and data analysis: AM, JN, ES, KS, MLP, LH, MW, JC, JM, JK, ZL, AW-T

Data science and machine learning: KS, JK, ZL, AW-T

Figures and data visualization: VL, KS, JK

Manuscript drafting: VL, CJ

Manuscript review and editing: all authors

## DATA AVAILABILITY

All processed molecular data (bulk RNA-Seq, whole genome sequencing, targeted mutation data) and corresponding scripts to generate figures are available on an S3 bucket (s3://deepcell_public/ mavropoulos_2022) and https://github.com/deepcell/mavropoulos_2022, respectively. Raw data can be provided upon reasonable request.

