## Supplemental Figures for "Artificial intelligence-driven morphology-based enrichment of malignant cells from body fluid"

**Supplementary Figure 1**

A

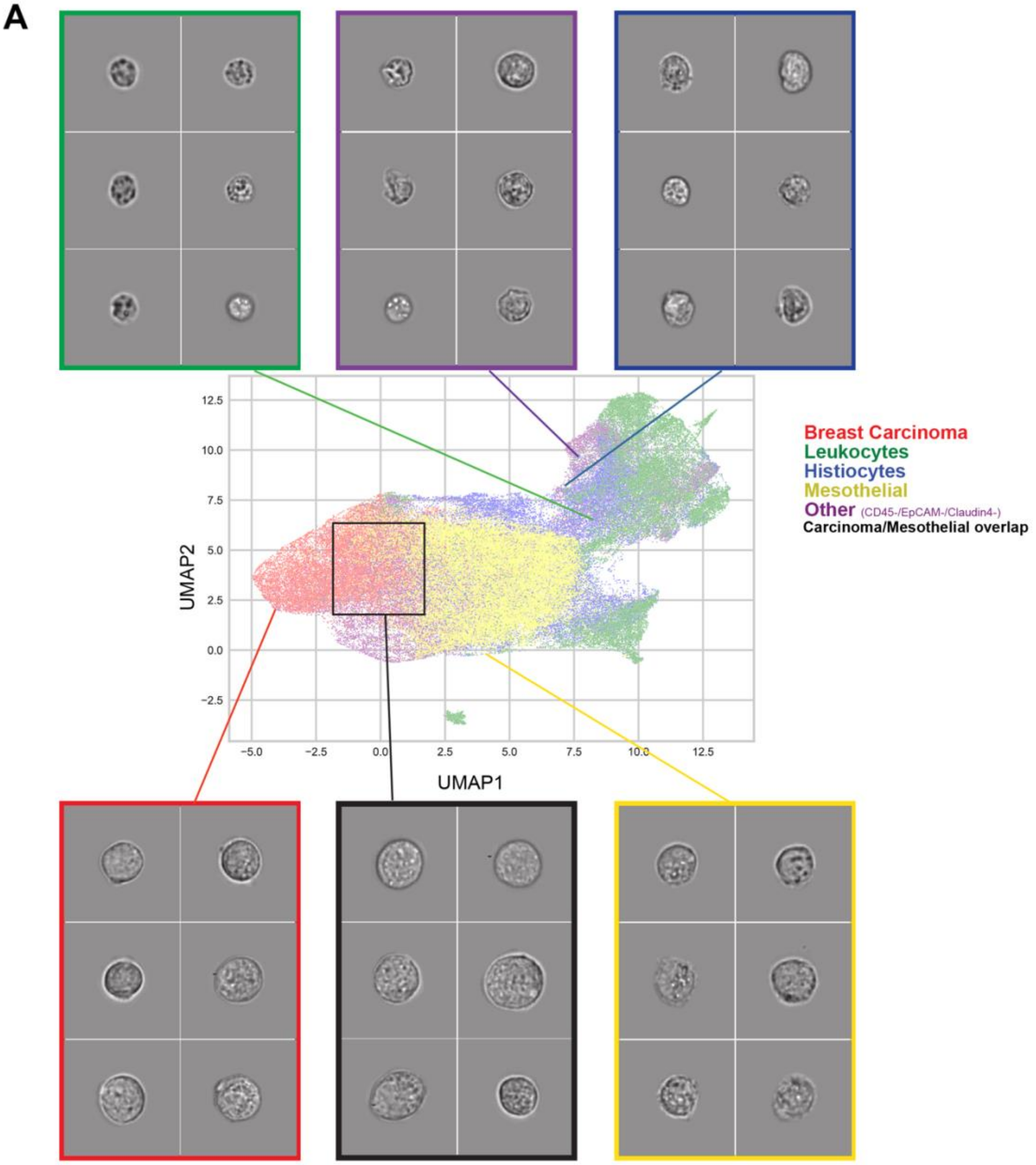


B

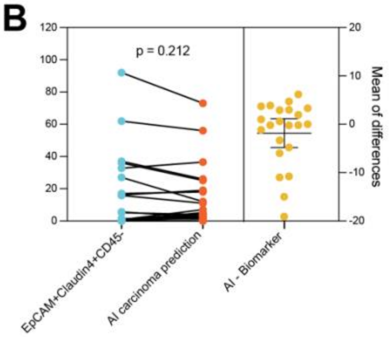


C

| Cancer type | EpCAM+Claudin4+CD45- | | | AI carcinoma prediction | | |
|---|---|---|---|---|---|---|
| | Mean | Median | Range | Mean | Median | Range |
| **Breast** (n=4) | 15.91% | 13.80% | 0.05-36.0% | 11.13% | 8.65% | 2.00-25.2% |
| **GI** (n=5) | 1.53% | 0.83% | 0.00-5.19% | 2.46% | 1.26% | 0.00-7.00% |
| **Gyn** (n=10) | 27.42% | 16.55% | 0.01-92.1% | 24.84% | 18.50% | 1.90-73.00% |
| **Lung** (n=3) | 2.30% | 1.08% | 0.21-5.61% | 2.23% | 2.30% | 0.00-4.39% |

**Supplementary Figure 1: Related to Figure 1**

**(A)** Embeddings UMAP plot of *in silico* mixture of known isolated cell types, related to **Fig. 1C**. Embeddings are extracted from images of the model validation dataset (15,000 images randomly sampled from >2 million validation set images). Coloring is based on known cell type (ground truth) and clustering patterns are based on morphological similarity detected by the AI model. Cells (dots) clustering closer from each other are more morphologically similar. Cell types are annotated based on cell lines or FACS-sorted pure population labels (see Methods). Breast carcinoma (red), leukocytes (green), histiocytes (blue), mesothelial (yellow), biomarker-negative cell types "other" (purple), and carcinoma/mesothelial cell overlap region (black). Representative brightfield images from each cluster are shown; for display only, these images have been normalized for illumination intensity, and the pixels outside the cell have been blanked out.

**(B)** Relative differences between proportion in percent of EpCAM+Claudin4+CD45- carcinoma cells (quantified by flow cytometry) and the proportion in percent of AI model carcinoma cell prediction (quantified by AI) in effusion samples. Paired t-test was performed, $p = 0.1212$. Related to **Fig. 1E-F**.

**(C)** Descriptive statistics of AI model and biomarker estimation values. Related to **Fig. 1E-F**.

**Supplementary Figure 2**

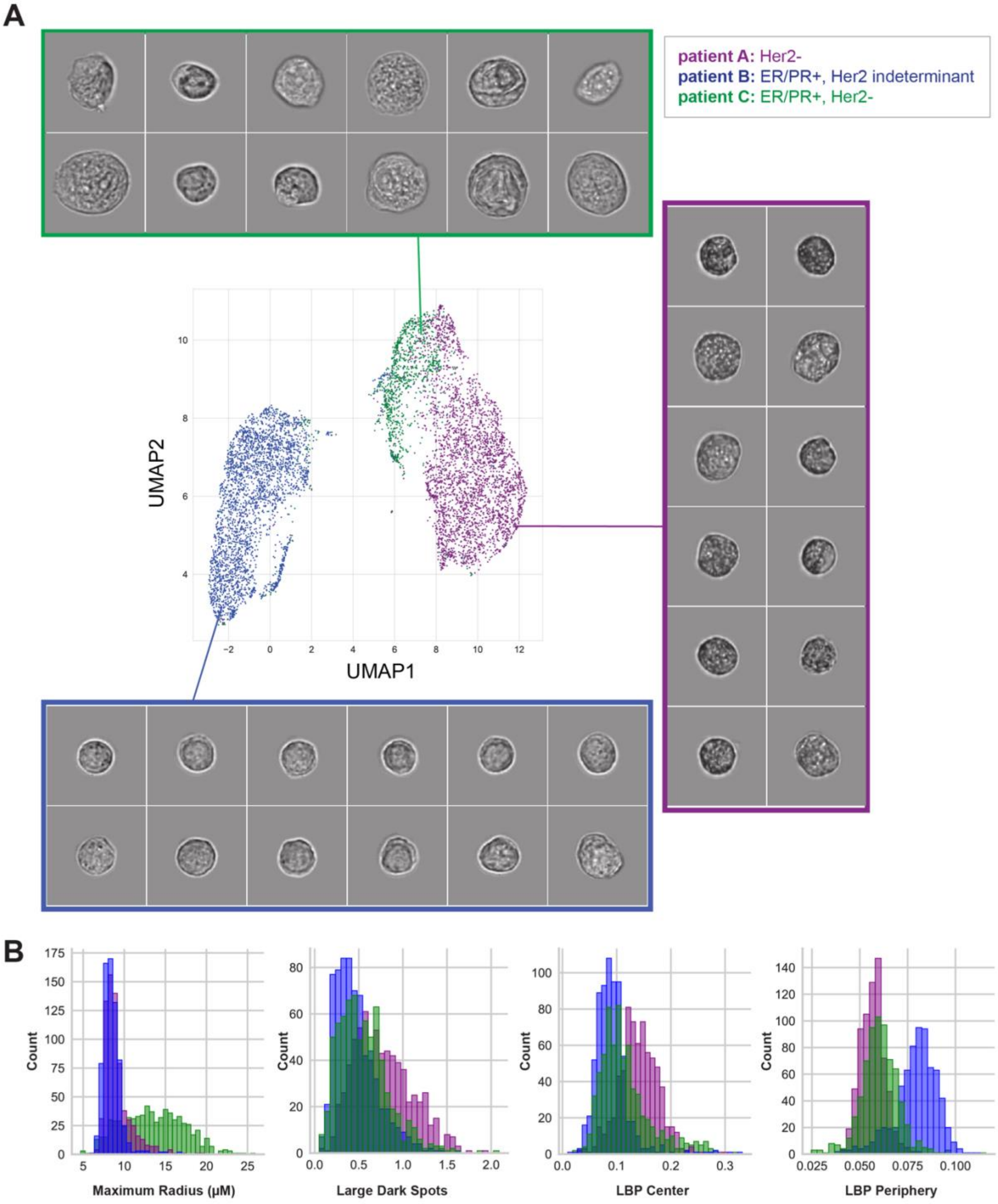

**Supplementary Figure 2: Related to Figure 2**

**(A)** Embeddings UMAP plot of AI prediction of carcinoma cells, as classified by the malignant effusion AI model. Embeddings are extracted from images of unlabeled breast cancer effusion samples from three different patients. Cancer subtype is indicated in the legend. Representative brightfield images from each cluster are shown, for display only.

**(B)** Histogram plot by patient, as quantified by computer vision morphometric measures of cell size (radius) and texture (large dark spots identified by a Laplacian of Gaussian filter, Local Binary Pattern (LBP) center, LBP periphery). LBP can efficiently describe texture in 10 numbers; the presented data shows the distribution of values of one of these numbers for the three different patients

**Supplementary Figure 3**

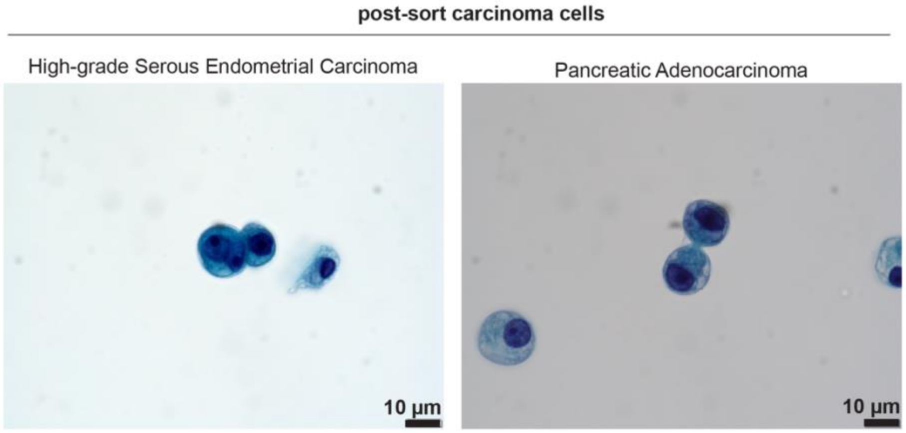

**Supplementary Figure 3. Related to Figure 3**

PAP stain of post-sort carcinoma cells enriched from high-grade serous endometrial carcinoma and pancreatic adenocarcinoma effusion samples. Scale bar as indicated.

**Supplementary Figure 4**

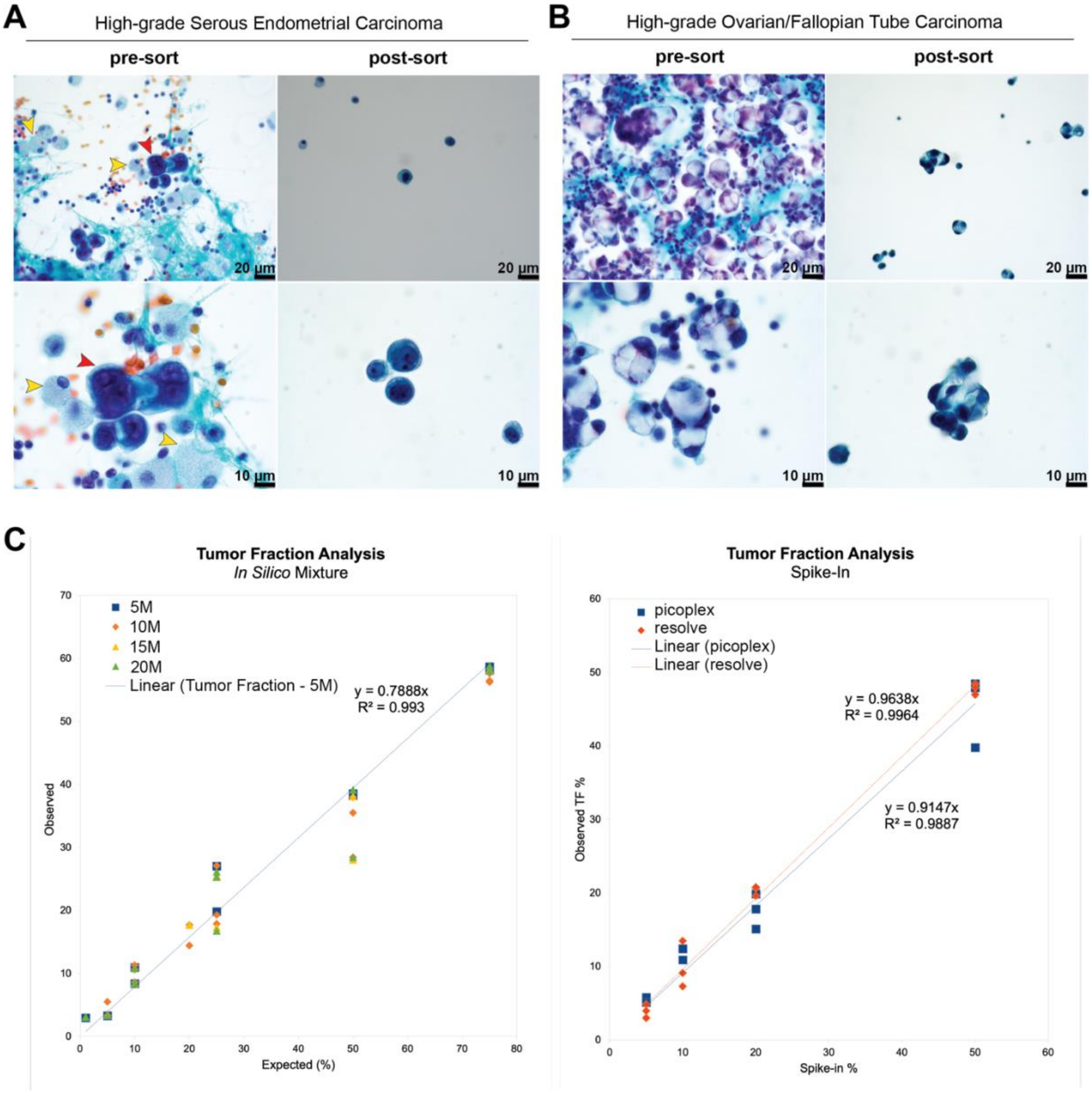

**Supplementary Figure 4. Related to Figure 4.**

**(A-B)** Additional PAP stains performed and assessed by a board-certified pathologist on pre- and post-sort **(A)** high-grade serous endometrial and **(B)** high-grade ovarian/fallopian tube carcinoma ascitic fluids at different magnifications. Red arrows indicate cells of interest/atypical cells, yellow arrow indicate inflammatory or histiocyte cells. Scale bar as indicated.

**(C)** Tumor fraction analysis is performed with the ichorCNA tool, both *in-silico* (left panel) and spike-in (right panel) data were used to confirm reported tumor fraction values.
